# Polymerase mutations underlie early adaptation of H5N1 influenza virus to dairy cattle and other mammals

**DOI:** 10.1101/2025.01.06.631435

**Authors:** Vidhi Dholakia, Jessica L. Quantrill, Samuel Richardson, Nunticha Pankaew, Maryn D. Brown, Jiayun Yang, Fernando Capelastegui, Tereza Masonou, Katie-Marie Case, Jila Ajeian, Maximillian N.J. Woodall, Callum Magill, Graham Freimanis, Amy McCarron, Ecco Staller, Carol M. Sheppard, Ian H. Brown, Pablo R. Murcia, Claire M. Smith, Munir Iqbal, Paul Digard, Wendy S. Barclay, Rute M. Pinto, Thomas P. Peacock, Daniel H. Goldhill

## Abstract

In early 2024, an unprecedented outbreak of H5N1 high pathogenicity avian influenza was detected in dairy cattle in the USA^1^. As of mid-2025 the epidemic is ongoing, resulting in spillbacks into poultry, wild birds and other mammals including humans^2^. Here, we present molecular and virological evidence that the cattle B3.13 genotype H5N1 viruses rapidly accumulated adaptations in polymerase genes that enabled better replication in bovine cells and tissues, as well as cells of other mammalian species including humans and pigs. We find evidence of several mammalian adaptations gained early in the evolution of these viruses in cattle including PB2 M631L, which is found in all cattle sequences, and PA K497R, which is found in the majority. Structurally, PB2 M631L maps to the polymerase-ANP32 interface, an essential host factor for viral genome replication. We show that this mutation adapts the polymerase to better interact with bovine ANP32 proteins, particularly ANP32A, and thereby enhances virus replication in bovine mammary systems and primary human airway cultures. Importantly, we show that ongoing evolution during 2024 and 2025 in the PB2 gene, including E627K and a convergently arising D740N substitution, further increase polymerase activity and virus replication in a range of mammalian cells. Thus, the continued circulation of H5N1 in dairy cattle not only allows virus adaption improving replicative ability in cattle, but also increases the risk of zoonotic spillover.

## Introduction

The natural host reservoirs of influenza A viruses (IAV) are wild waterfowl and shorebirds. However, IAV can spillover and adapt to infect mammalian species, such as pigs, humans, horses and dogs^3^. In 2020, a novel genotype of H5N1 clade 2.3.4.4b emerged in Europe and spread around the world causing a panzootic in birds^4^. This panzootic genotype and its progeny are notable for their frequent spillover into mammals, including confirmed or potential mammal-to-mammal spread^5^ in farmed mink in Europe^6–8^ (2022-2023), semi-aquatic marine mammals in South America and Antarctica^9–13^ (2023-), and recently, dairy cattle in the USA^1,14^ (2024-).

In early 2024, an unexplained drop in milk production in cattle in the USA was attributed to infection with H5N1 clade 2.3.4.4b, of the B3.13 genotype^1,14^. Infected cows were initially observed in Texas but as of June 2025, over 1000 infected herds have been recorded across 17 US states^1,14^. This outbreak was most likely caused by a single spillover from wild birds^1,14,15^, with continuous transmission between dairy cattle (hereafter referred to as ‘cattle’) mediated by human activities such as contamination of milking machinery within farms^16^, and movement of infected animals and/or contaminated equipment between farms and states^1^. The virus does not cause high mortality in cattle, instead mostly causing mastitis and modest respiratory distress^14,17^. In addition to cattle, several other species have been infected following direct or indirect exposure to infected cattle including chickens, turkeys, peridomestic birds, farmed alpaca, domestic cats^1^, and humans^2,18^. Infections have been repeatedly fatal in cats that ingested infected milk, but in humans the B3.13 virus has been associated with conjunctivitis and/or mild respiratory disease^2,18^.

Avian influenza viruses (AIVs) usually have restricted replication in mammalian hosts^3^. However, they can rapidly accumulate mammalian adaptations that overcome this block, often in the viral RNA-dependent RNA polymerase^5,19,20^. In particular, ANP32 proteins are essential host-encoded factors co-opted by the virus to support genome replication^21,22^. All birds excluding palaeognaths (e.g. ostriches and related birds), have an ANP32A gene with an additional exon resulting in splice variants encoding proteins with an extra 33 amino acids relative to mammalian orthologues^23^. Avian-origin influenza viruses require polymerase mutations to utilise the shorter mammalian ANP32A/B proteins^24^. The most widespread, and best characterised adaptive mutations are those in the PB2 subunit of the polymerase including PB2 E627K^23^, Q591R/K^25,26^, and D701N^26,27^, all of which have been detected during other mammalian H5N1 outbreaks^7–13,28^ but are absent in the initial dairy cattle viruses. However, alternative adaptive mutations in the influenza polymerase that enable the virus to co-opt suboptimal ANP32 proteins are possible^29–31^. Here, we investigated potential mammalian adaptations in the polymerase of cattle-adapted H5N1 viruses and identify two key mutations that allow the B3.13 virus to replicate in cells and tissues from cattle and other mammalian species, including humans.

## Results

### Phylogenetic analysis of the cattle H5N1 polymerase identified potential adaptive mutations

To understand how an avian influenza virus had adapted to efficiently replicate in cells of a novel mammalian species, we constructed a phylogeny using concatenated genomes of all influenza A viruses isolated from cattle, virus from the first reported human case and wild avian viruses from September 2023-March 2024 (Supplementary Figure S1). Viruses from cattle formed a monophyletic group suggesting a single spillover event from wild birds^1,15^ (Figure 1A). The sequence of the first case of human infection with a B3.13 H5N1 virus^2^ was an outlier relative to all subsequent cattle B3.13 viruses (Figure 1A, Supplementary Figure S1) and contained the known mammalian adaptations PB2 E627K^32^ as well as PA K142E^33^ relative to the closest avian relatives (Supplementary Figure S1). However, these adaptive mutations were not found in any cattle virus sequences, which instead all contained PB2 M631L as well as PB2 E362G and PA L219I. PB2 M631L has previously been reported to adapt avian H1N1, H5N1 and H10N7 viruses to mammals^32,34,35^, as well as H9N2 viruses to gene-edited chickens with a non-functional ANP32A^31^. Early cattle sequences grouped into three clades with additional mutations in the polymerase segments including i) PB2 E677G^36,37^, ii) PA I13V and E613K, and iii) PA K497R, which was found in the largest cattle clade (Figure 1A, Supplementary Figure S1), consisting of ∼95% of sequences (as of December 2024) and has previously been associated with influenza adaptation to mammals^38^. Minor clade i (PB2 E677G) was found solely in Texas and appeared only at the beginning of the outbreak whereas minor clade ii (PA I13V) was detected across multiple states, including, most recently, in Colorado.

**Figure 1.**
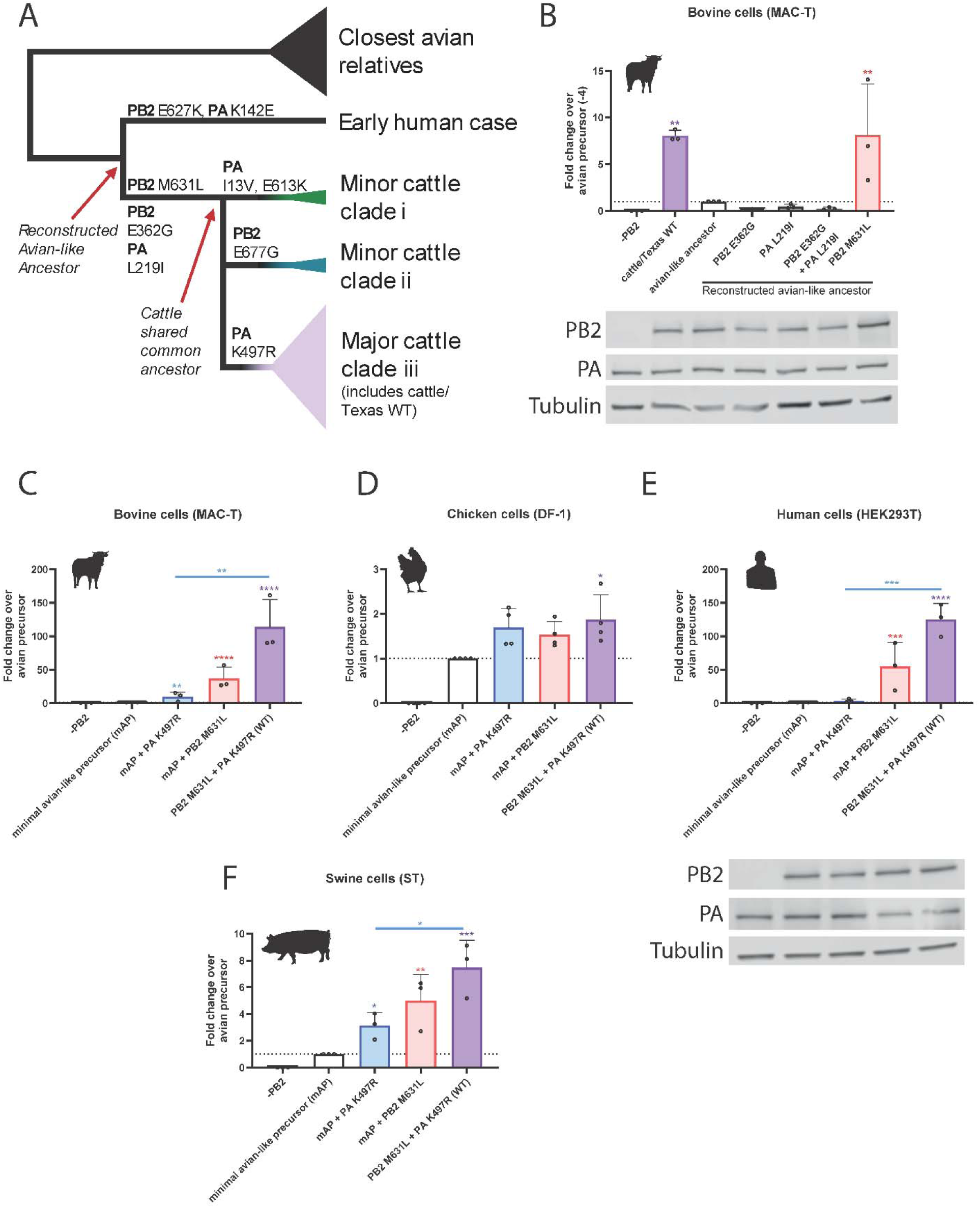
Identification of key mammalian polymerase adaptations in the major clade of bovine H5N1 that enhance polymerase activity in bovine cells. A) Simplified schematic of cattle polymerase phylogeny and polymerase mutations that arose in early branches. B-F) Minigenome in (B, C) bovine MAC-T cells (D) chicken DF-1 (E) human HEK293T, (F) swine ST cells with cattle Texas (WT). B) Reconstructed avian-like ancestor has a reversion of PB2 E362G, M631L and PA L219I, K497R. C-F) Minimal avian-like precursor has the reversions of PB2 M631L and PA K497R. Data normalised to minimal avian-like precursor. Data throughout plotted as the mean of N = 3 independent repeats. (B, E) Expression of PB2 and PA was measured by western blot, tubulin was used as a loading control. Data throughout plotted as mean + SD. Statistics throughout performed by one-way ANOVA with multiple comparisons comparing all data using log-transformed data. B) significance only plotted if increased over avian-like ancestor. Log-normality determined by Shapiro-Wilk test and QQ plot. Significance shown by asterisks indicating: *, 0.05 ≥ P > 0.01; **, 0.01 ≥ P > 0.001; ***, 0.001 ≥ P > 0.0001; ****, P ≤ 0.0001.

### Activity of bovine H5N1 polymerase in mammalian cells maps to PB2 M631L and PA K497R

To test whether the mutations identified from the phylogenetic analysis were adaptive in cattle, we first constructed a ‘minireplicon’ system suitable for testing AIV polymerase activity in bovine cells. This consisted of co-transfecting bovine cells with a custom-made minigenome reporter that uses a bovine RNA polymerase I (Pol I) promoter to express a viral-like RNA encoding firefly luciferase flanked with non-coding regions from influenza A segment 8/NS^39^ along with plasmids expressing influenza virus ribonucleoprotein components, PB2, PB1, PA and nucleoprotein (NP). This species-specific minireplicon system showed 10-20-fold higher activity in bovine cells than one using an equivalent human PolI reporter (Supplementary Figure S2). Using this bovine-specific minireplicon reporter, we showed that the reconstituted polymerase complex of A/dairy cattle/Texas/24-008749-001-original/2024 (cattle/Texas) was highly active in MAC-T bovine mammary epithelial cells^40^ (Figure 1B).

We next reconstructed an avian-like ancestor to represent the polymerase complex of the virus that mostly likely entered the cattle population, by reverting potential adaptive mutations from the PB2 and PA segments of cattle/Texas: PB2 M631L, PB2 E362G, PA K497R and PA L219I (Figure 1A, Supplementary Figure S1). Whilst still active above background, the avian-like ancestral polymerase was much less active than its WT cattle/Texas counterpart (Figure 1B). To begin to identify key changes, we assessed the impact of adding back the mutations that arose between this reconstructed avian-like ancestor, and the cattle virus shared common ancestor (PB2 M631L, PB2 E362G and PA L219I). Neither PB2 E362G nor PA L219I increased polymerase activity in bovine cells relative to the avian-like ancestor background, either individually or in combination, and PB2 E362G even reduced activity (Figure 1B). In contrast, PB2 M631L increased activity by around 10-fold, suggesting this was the key polymerase mutation that arose early in the adaptation of H5N1 to dairy cattle.

We also investigated later mutations that arose in the major clade iii of the cattle H5N1 virus (Figure 1A). We first tested the effect of the PA mutation K497R that is maintained across all sequences of the major bovine clade. Reverting both PB2 631L to M and PA 497R to K (generating a ‘minimal avian-like precursor’ or mAP) resulted in a ∼100-fold decrease in polymerase activity in bovine cells (Figure 1C). The reintroduction of PB2 M631L alone increased the signal by around 50-fold. Reintroducing PA K497R had a more modest effect (∼10-fold increase in signal). Neither mutation alone fully recapitulated the activity of the WT polymerase, indicating that the PA change was further adaptive. The same mutations had minimal impacts (<2-fold) in avian cells (chicken fibroblasts, DF-1; Figure 1D), consistent with an adaptive function in cattle cells rather than simply altering the intrinsic activity of the viral polymerase.

We next tested the impact of these two mutations on polymerase activity in cells from two other mammalian hosts relevant to influenza viruses - humans and swine, again using host species-specific minireplicon reporters (Figure 1E, F). As seen in the bovine cells, both adaptations individually significantly enhanced polymerase activity, with PB2 M631L having the greater effect, suggesting it is the major mammalian adaptation across different mammalian species. Also, as observed in bovine cells, we noted a cumulative effect of the two substitutions together.

To investigate whether PB2 M631L and PA K497R could be universal mammalian adaptations for H5 viruses, we introduced them into other H5N1 polymerase backgrounds: a recent 2.3.4.4b virus (AIV07-A/chicken/England/053052/2021), an EU reference laboratory (EURL) genotype C common in the UK in 2021/22 and a distantly related H5N1 IAV polymerase (A/turkey/England/50-92/1991; 50-92), both of which originated from low pathogenicity avian influenza virus donors^41^. Relative to cattle/Texas, AIV07 and 50-92 have amino acid identities for PB2 of 98% and for PA of 98% and 97%, respectively. In these additional genetic backgrounds, both PB2 M631L and PA K497R significantly enhanced polymerase activity in bovine cells and human cells (Supplementary Figure S3A-F). When compared to the well-characterised mammalian adaptation PB2 E627K, we saw that neither individual mutation increased polymerase activity to the level of E627K, but when combined, there was no statistical difference seen between PB2 M631L/PA K497R and E627K, suggesting the combined mutations are just as potent. There was, however, a trend that E627K alone had a larger effect relative to PB2 M631L + PA K497R in the human cells compared to the cattle cells. Overall, this suggests both PB2 M631L and PA K497R are *bona fide* mammalian adaptations, relevant across diverse AIVs.

### PB2 M631L, and to a lesser extent PA K497R, enable efficient replication of bovine H5N1 in mammalian cells and explants

To further test the importance of PB2 M631L and PA K497R in adapting virus to replicate in mammalian cells, we generated a reverse genetics system for cattle/Texas and made viruses bearing mutations at these sites. To work safely at BSL2, we rescued viruses as 2:6 reassortants containing the HA and NA of the attenuated vaccine strain, A/Puerto Rico/8/1934 (PR8)^42^ and the remaining genes from cattle/Texas. We also conducted work with full reverse genetics derived viruses at ACDP3/SAPO4 (BSL3+). We compared multicycle replication of the WT cattle/Texas virus to either the individual or double PB2/PA reversions in bovine *ex vivo* mammary gland explants (Figure 2A). Compared to the minimal avian-like precursor (WT cattle/Texas virus with PB2 M631L and PA K497R reverted), the viruses with PB2 M631L alone or in combination with PA K497R showed a significant, >10-fold increase in infectious virus titres at 24 and 48 hours. However, PB2 M631L alone grew to significantly lower titres than the WT virus that contained K497R as well at 48 and 72 hours post-infection (Figure 2A). This was reflected both in released virus from the explants, as well as tissue associated virus harvested at 72 hours post-infection (Figure 2B).

**Figure 2.**
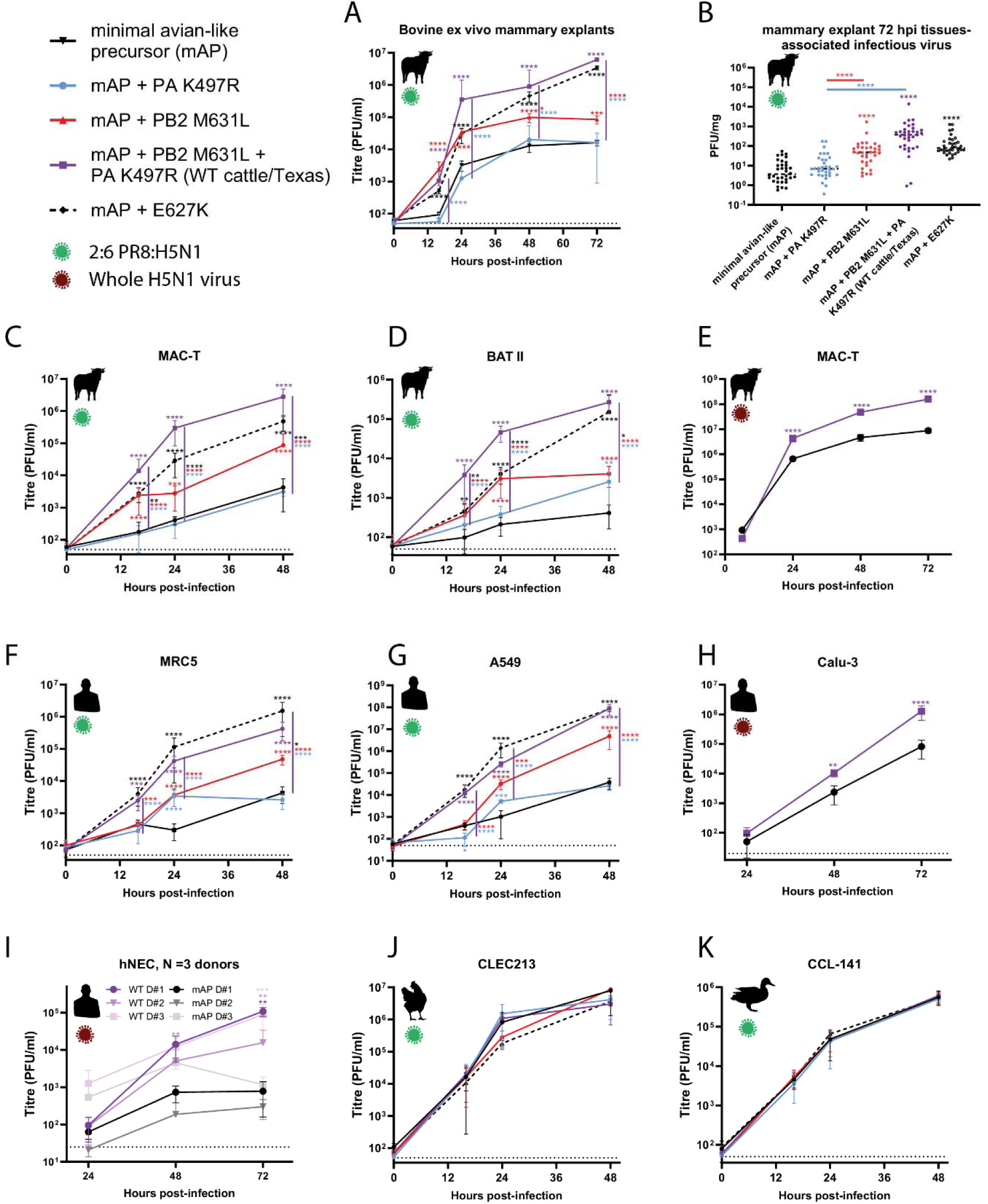
PB2 M631L, and to a lesser extent PA K497R, enhance virus replication in mammalian, but not avian culture systems. Reverse-genetics derived viruses, containing the HA and NA genes of the attenuated laboratory strain A/Puerto Rico/8/34, and the remaining genes from Cattle Texas (2:6 viruses; panels A-D, F,G,J,K) or reverse genetics derived full H5N1 viruses (whole virus, panels E, H and I) were used to infect A, B) bovine ex vivo mammary explants, C, E) bovine mammary cells (MAC-T), D) bovine respiratory cells (BAT II) cells, F) human lung cells (MRC5), G) human lung adenocarcinoma cells (A549), H) human lung cells (Calu-3), I) primary human nasal epithelial cultures (hNECs) maintained at air liquid interface. J) chicken lung cells (CLEC213) or K) duck fibroblasts (CCL-441). Explants were infected with 5000 pfu/explant. Cell lines were infected at an MOI of 0.01. Infectious virus titres were determined by plaque assay on MDCKs. Data from bovine explants represents N = 34 repeats from 3 donor cattle from 3 different independent experiments. Data from cell lines plotted as (C), 2 x N = 4, (D,F,G,J, K) 2 x N = 3 technical repeats, (E,H) N = 3 repeats, from a representative repeat of N = 3 independent repeats and I) 3 independent donors with virus replication tested in triplicate. Data throughout plotted as mean ± SD. Statistics throughout performed by two-way ANOVA on log-transformed data with multiple comparisons between all data points within a single time point. Significance only plotted when differences were significant to the minimal avian precursor (shown as matched colour asterisks), or between the WT cattle/Texas and mutants (shown as coloured asterisks next to the purple line). Significance shown by asterisks indicating: *, 0.05 ≥ P > 0.01; **, 0.01 ≥ P > 0.001; ***, 0.001 ≥ P > 0.0001; ****, P ≤ 0.0001.

Consistent results were also seen in MAC-T cells and bovine alveolar cells (BAT II; Figure 2C-E) where PB2 M631L, but not PA K497R alone significantly enhanced replication across multiple tested time points. Again, PB2 M631L replicated to significantly lower titres than the wild type virus also containing PA K497R. Overall, these results in the context of infectious virus are consistent with the minireplicon data that show PB2 M631L, and to a lesser degree PA K497R, are responsible for this virus’s ability to replicate in bovine cells.

We next tested the replication of these viruses in several human lung cell lines (MRC5, A549 and Calu3; Figure 2F-H). Consistent with the minireplicon assays in human cells (Figure 1E) and virus replication phenotypes in bovine cells (Figure 2A-E), the PB2 M631L mutation alone, or combined with PA K497R, significantly enhanced virus titres compared to the minimal avian-like precursor virus across multiple time points in the human cell lines. While PA K497R alone showed only minor effects, PB2 M631L alone replicated to significantly lower titres than the WT virus (cattle/Texas), again showing PA K497R, in particular, boosted replication when paired with PB2 M631L. We also tested the replication of the WT and mAP virus in primary human nasal epithelial cultures (hNECs) from 3 adult donors and maintained at air liquid interface (Figure 2I). The mAP virus replicated to significantly lower titres than WT in all 3 donors at 72 hours post-infection. Overall, these data further establish that PB2 M631L is a major mammalian adaptation that enhances virus replication in both bovine and human cells.

Next, we tested the growth of these viruses in a pair of avian cell lines (chicken CLEC213 and duck CCL-141; Figure 2J,K). All mutants showed similar growth kinetics to the wild type virus, suggesting that these mutations have a minimal impact on virus fitness/replication in avian cells.

Across the growth experiments in Figure 2, we also compared the relative replication of the mAP and mAP containing PB2 E627K. Consistently, across all bovine cell lines and explants (Figure 2A-D), the combination of PB2 M631L and PA K497R resulted in significantly higher titres than mAP + E627K. Conversely, in the human cells (Figure 2F,G), the mAP + E627K virus replicated to higher titres (albeit not significantly) than the WT cattle virus. Overall, this might suggest why PB2 M631L and PA K497R were favoured in the cattle virus, whereas PB2 E627K is more common in human infections.

All available H5N1 sequences from this cattle outbreak contain PB2 M631L but PA K497R is only found in the major clade of cattle sequences (clade iii). Minor clade i and ii viruses contain alternative PB2 or PA mutations (Figure 1A, supplementary Figure 1). To investigate whether these minor clade mutations represent alternative bovine adaptations, we recapitulated these genotypes by individually introducing PA I13V and E613K from clade i or PB2 E677G from clade ii into the cattle/Texas polymerase background with PA R497 reverted to the avian consensus 497K. PA I13V and E613K together exerted a small but significant increase in polymerase activity over PB2 M631L alone showing a ∼3-fold increased minireplicon signal in bovine cells (Supplementary Figure S4A) and human cells (Supplementary Figure S4B). The introduction of PB2 E677G had no impact in bovine cells but resulted in a significant ∼4-fold increase in polymerase activity in human cells (Supplementary Figure S4B). The polymerase mutations found in the minor clade i and ii cattle viruses had at most a very minor impact (<1.5-fold) in polymerase activity in avian cells (Supplementary Figure S4C). Overall, this suggests that PB2 M631L was the initial potent mammalian adaptation, and a range of other polymerase mutations then ‘fine-tuned’ polymerase activity in bovine cells.

### More recent mutations in cattle sublineages further enhance B3.13 H5N1 polymerase activity in mammalian but not avian cells

By continually monitoring sequences from the ongoing dairy cattle outbreak, we detected incidences of further, known mammalian adaptations arising within polymerase genes of the cattle viruses^26^. This included a single emergence of a cluster of sequences containing PB2 Q591R, at least three independent emergences of PB2 D740N across the tree (Supplementary Figure S1), as well as a more recent cluster from 2025 containing PB2 E627K. To test the impact of these changes we introduced them into the WT cattle/Texas polymerase. PB2 D740N significantly boosted polymerase activity in bovine (∼4-fold increased signal), human (∼2-fold increase signal) and swine cells (1.2-fold increase; Figure 3A-C). PB2 Q591R led to a smaller impact, reaching statistical significance in the human cells (Figure 3A, B). The introduction of PB2 E627K on top of M631L consistently led to the highest polymerase activity (Figure 3A-C). However, neither PB2 E627K nor D740N increased activity in avian cells, relative to the wild-type cattle/Texas polymerase (Figure 3D). We conclude that these mutations indicate ongoing adaptation of H5N1 viruses to further optimise polymerase function in cattle.

**Figure 3.**
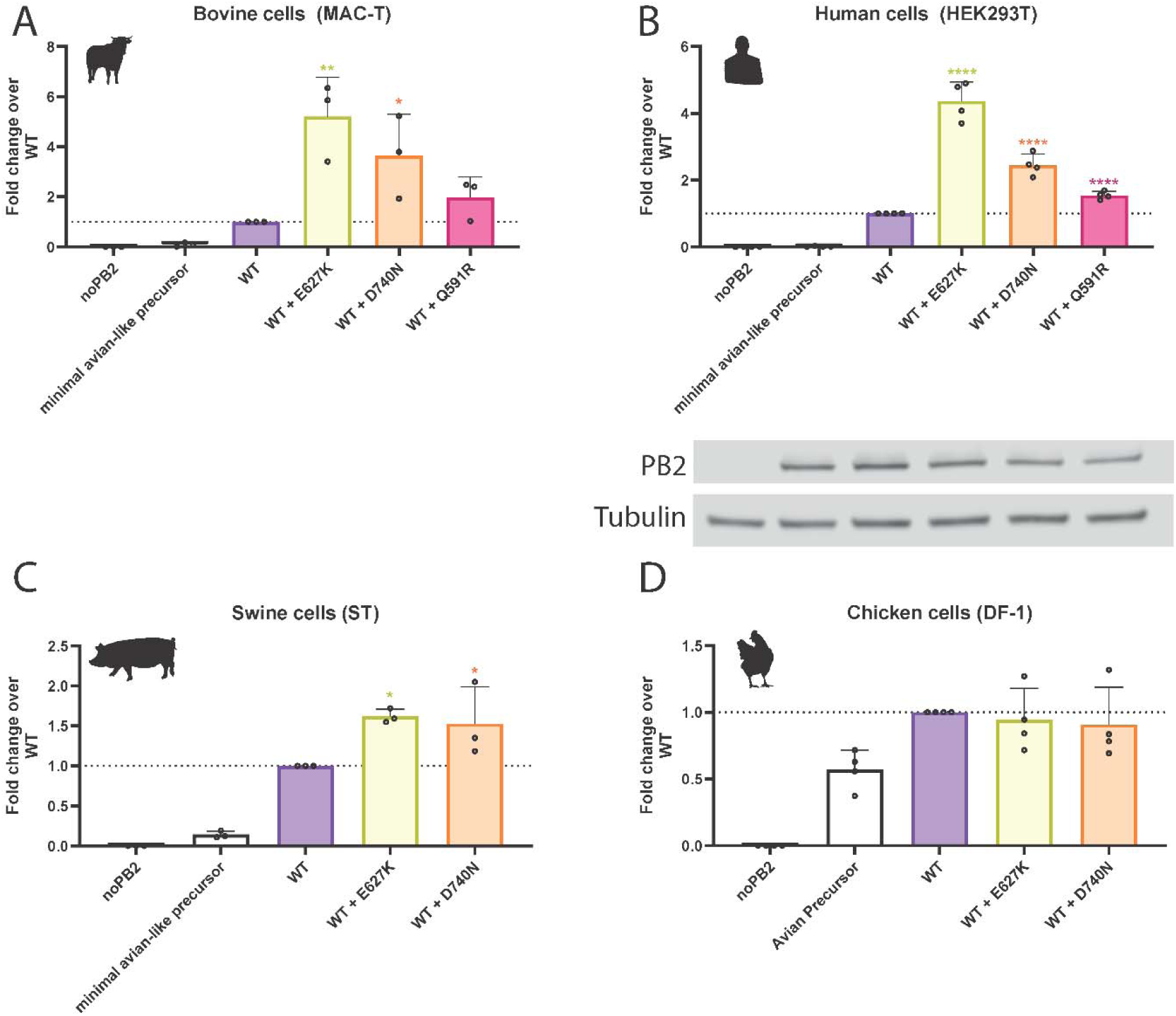
Ongoing adaptations found in cattle H5N1 viruses further enhance polymerase activity in mammalian cells. Minigenome in (A) bovine MAC-T cells, (B) human HEK293T, (C) swine ST cells or (D) chicken DF-1 with mutant versions of cattle Texas representing potential ongoing adaptation. Data normalised to WT cattle Texas. Data throughout plotted as mean of N = 3 independent biological repeats. B) Matched western blot showing PB2 expression. Data throughout plotted as mean + SD. Statistics throughout performed by one-way ANOVA with multiple comparisons against WT polymerase using log-transformed data. Log-normality determined by Shapiro-Wilk test and QQ plot. Significance shown by asterisks indicating: *, 0.05 ≥ P > 0.01; **, 0.01 ≥ P > 0.001; ***, 0.001 ≥ P > 0.0001; ****, P ≤ 0.0001.

To further investigate the impact of these ongoing mammalian adaptations, we rescued WT cattle/Texas + PB2 D740N. For work at CL2, we also rescued PR8:H5N1 2:6 reassortants of cattle/Texas with PB2 D740N or PB2 E627K. We initially tested the ability of the viruses to replicate in bovine mammary explants, and found that while PB2 D740N closely tracked the wild type virus, PB2 E627K reached significantly higher titres at early time points (Figure 4A). Similar results were seen in MAC-T and BAT-II cells where both D740N and E627K significantly enhanced replication in the WT across multiple time points (Figure 4B,C).

**Figure 4.**
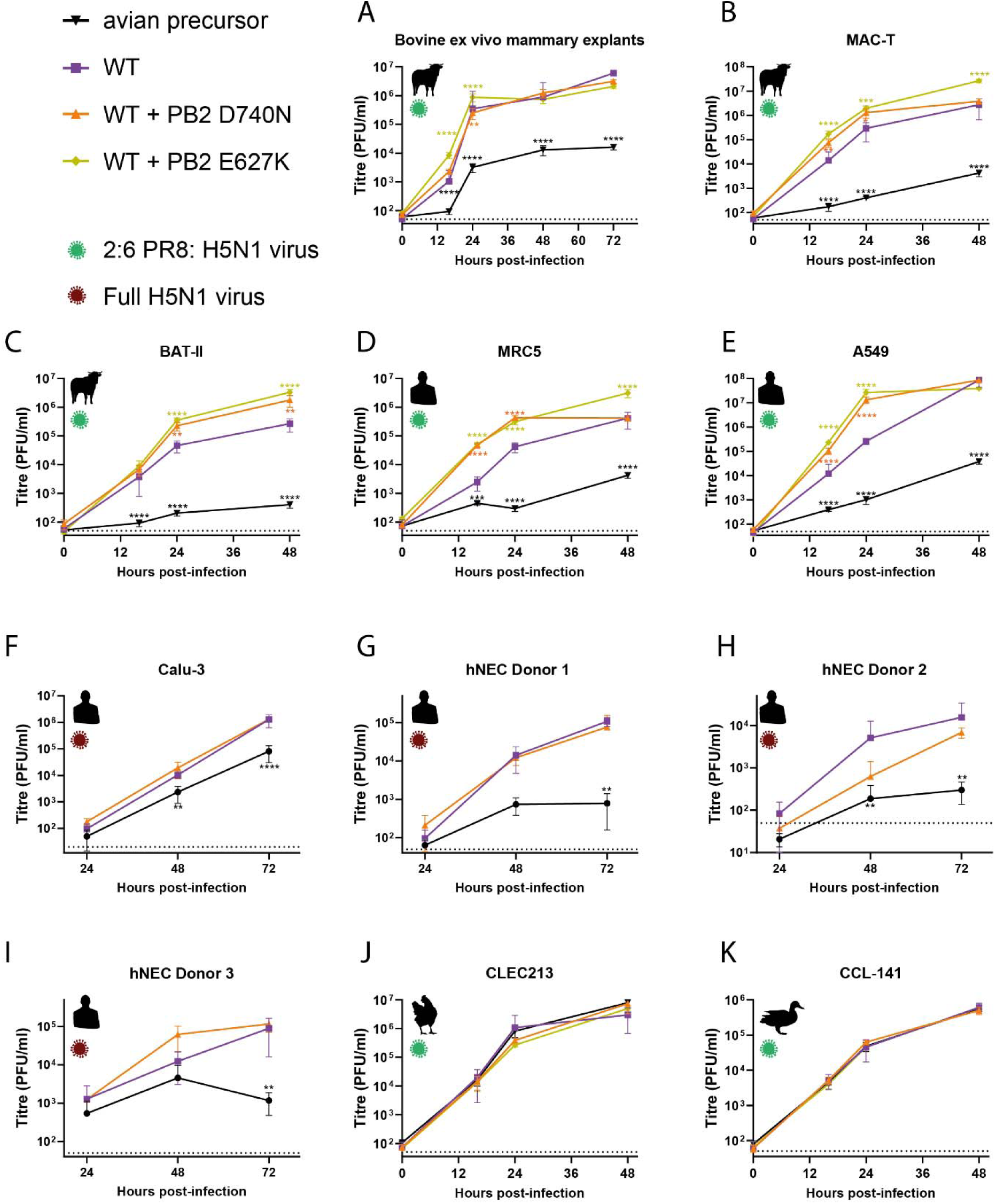
Further emerging mammalian adaptations in the cattle H5N1 PB2 enhance its replication in human and bovine, but not avian culture systems. Reverse-genetics derived viruses, containing the HA and NA genes of the attenuated laboratory strain A/Puerto Rico/8/34, and the remaining genes from Cattle Texas (2:6 viruses; panels, A-E, J, K) or reverse genetics derived full H5N1 viruses (whole virus, panels F-I) were used to infect) A bovine *ex vivo* mammary explants, B) bovine mammary cells (MAC-T), C) bovine respiratory cells (BAT II) cells, D) human lung cells (MRC5), E) human lung adenocarcinoma cells (A549), F) human lung cells (Calu-3), G-I) primary differentiated human nasal epithelial cultures (hNECs) maintained at air liquid interface (hNECs), J) chicken lung cells, or K) duck fibroblasts (CCL-141). Cells were infected with an MOI of 0.01, explants were infected with 5000 PFU/explant. Infectious virus titres were determined by plaque assay on MDCKs. A) Data from bovine explants represents total N = 34 repeats from 3 donor cattle from 3 different independent experiments. Data from cell lines plotted as (B), 2 x N = 4, (C-E,J,F) 2 x N = 3 technical repeats, (F) N = 3 repeats, from a representative repeat of N = 3 independent repeats and G-I) 3 independent donors with virus replication tested in triplicate. WT and avian precursor data through the figure is the same shown through Figure 2. Data throughout plotted as mean ± SD. Statistics throughout performed by two-way ANOVA on log-transformed data with multiple comparisons against WT cattle/Texas. Significance shown by asterisks indicating: *, 0.05 ≥ P > 0.01; **, 0.01 ≥ P > 0.001; ***, 0.001 ≥ P > 0.0001; ****, P ≤ 0.0001.

We also tested 2:6 virus in MRC5 and A549 cells (Figure 4D,E) and found both D740N and E627K significantly increased virus titres across multiple time points. In a human lung cell line (Calu-3), using the full H5N1 virus, D740N trended towards higher, albeit but non-significantly higher titres than WT virus (Figure 4F). Finally, we performed multicycle replication curves with whole H5N1 virus in primary differentiated human nasal epithelial cultures (hNECs), maintained at air liquid interface, taken from three independent adult donors (Figure 4G-I). PB2 D740N behaved variably between donors and did not reach titres significantly differently to the WT virus. We performed whole genome sequencing on the H5N1 viruses from two of the donors at 72 hours post-infection and found no evidence of adaptation or selection at the consensus level following virus replication.

Finally, we tested virus replication kinetics (in the 2:6 background) in chicken and duck cells (Figure 4J, K). We saw no significant difference in virus replication kinetics.

The PB2 E627K virus showed consistently higher titres than the wild type suggesting this mutation likely increases the ability to replicate in both human and bovine cells, while the PB2 D740N containing viruses showed a variable phenotype, suggesting that if this mutation does impact the virus, the effect is subtle and not consistently captured by these assays. Both E627K and D740N also appear to have a neutral impact on the ability to replicate in avian cells, suggesting these mutations could be maintained if viruses spill back into avian species.

### Mutations in the polymerase of the cattle H5N1 viruses adapt the polymerase to use bovine ANP32 proteins

PB2 residue 631 is located close to the interface between influenza polymerase and the host factor ANP32 (Figure 5A, Supplementary Figures S5), nearby several other mammalian ANP32-adapting mutations such as PB2 E627K and Q591R^26,43^. We therefore hypothesised that the B3.13 polymerase adapted to bovine cells by increasing its ability to co-opt bovine ANP32 proteins. To test this, we performed minigenome assays in human cells which lack ANP32A, ANP32B and ANP32E (eHAP triple knockout cells^30^) complemented with ANP32 proteins from other relevant species including cattle, human, pigs, and chicken. Only chicken ANP32A robustly supported viral polymerases from avian (AIV07, an early European 2.3.4.4b panzootic genotype C virus; Figure 5B) or avian-like viruses (the minimal avian-like precursor; Figure 5C). Neither human or bovine ANP32A or B supported any significant activity of avian virus polymerases. In contrast, both human and bovine ANP32A/B supported mammalian-adapted polymerases: cattle/Texas (Figure 5D) and a pandemic 2009 H1N1 virus (Figure 5E), A/England/195/2009 (Eng195). However, bovine ANP32A was consistently and significantly better able than bovine ANP32B to support polymerase activity of all constellations tested. We also assessed the effect of the individual cattle/Texas mammalian adaptations, PB2 M631L and PA K497R on complementation by the different ANP32 homologues (Figure 5F). Either mutation alone significantly enhanced polymerase activity supported by bovine ANP32A or ANP32B, with PB2 M631L showing a stronger effect with all ANP32 orthologues than PA K497R. Thus, we conclude that the major adaptive mutation PB2 M631L detected in the bovine influenza outbreak has adapted IAV for replication in cattle by increasing its capacity to co-opt bovine ANP32A.

**Figure 5.**
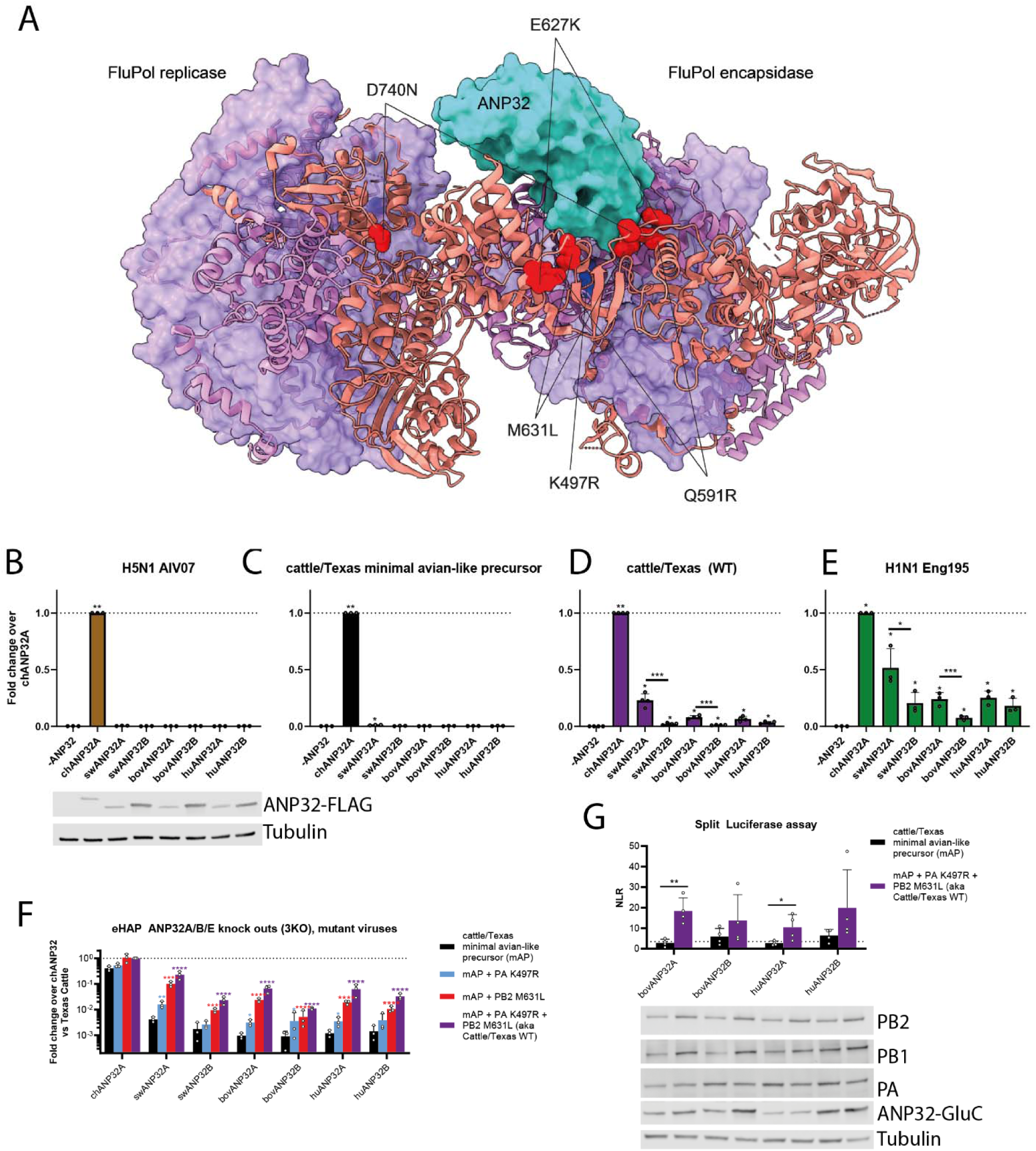
Although bovine ANP32A and B proteins are both able to support mammalian-adapted polymerases, bovine ANP32A is dominant which correlates with a stronger interaction. A) Key mammalian adaptations identified in this study mapped to the structure of H5N1 influenza A replicase dimer complex with human ANP32B (PDB: 8R1J). ANP32B is shown in cyan, PB1 in purple, PA in violet and PB2 as a salmon ribbon representation. H5N1 cattle PB2 substitutions (Q591R, E627K, M631L and D740N) shown in red and PA (K497R) shown in blue. B-F) Minigenomes in human engineered haploid cells (eHAP) with endogenous ANP32A, ANP32B and ANP32E knocked out were supplemented with ANP32 proteins from different species (chANP32A = chicken ANP32A, swANP32A = swine ANP32A, swANP32B = swine ANP32B, bovANP32A = bovine ANP32A, bovANP32B = bovine ANP32B, huANP32A = human ANP32A, huANP32B = human ANP32B, -ANP = no ANP32 control). Avianised cattle = PB2 M631L and PA K497R reverted. Data normalised throughout to chANP32A. G) Split luciferase assays in 293T cells showing the relative binding of different ANP32 proteins to the trimeric polymerase from cattle/Texas or the avianised version lacking PB2 M631L and PA K497R. PB1 was tagged with the N-terminal section of *Gaussia* luciferase, while the ANP32 proteins were tagged with the C-terminal portion. NLR, normalised luminescence ratios, were calculated from the ratio between signals between tagged and untagged ANP32/PB1 pairs. Dotted line equals a false positive cutoff of <2.5% as used in Mistry et al^61^. B, G) Matched expression of polymerase and ANP32 constructs shown by western blot. Data throughout plotted the mean of N = 3 (B-F) or N = 4 (G) independent biological repeats. Data throughout plotted as mean + SD. B-E) Statistics performed by one-way ANOVA with multiple comparisons - against -ANP32 control, or between species-matched ANP32A and ANP32B proteins, using log-transformed data. F) Statistics performed by two-way ANOVA with multiple comparisons comparing the different polymerase mutants within the same species ANP32, on log-transformed data. G) Statistics performed by multiple unpaired T tests between cattle/Texas WT and the minimal avian precursor. Log-normality determined by Shapiro-Wilk test and QQ plot. Significance shown by asterisks indicating: *, 0.05 ≥ P > 0.01; **, 0.01 ≥ P > 0.001; ***, 0.001 ≥ P > 0.0001; ****, P ≤ 0.0001.

Finally, to test the mechanism by which the bovine H5N1 virus was able to better use bovine ANP32 we performed a split-luciferase assay to test protein-protein interactions (Figure 5G). We tagged the cattle/Texas PB1 with one half of a split *Gaussia* luciferase and tagged different ANP32 proteins with the other half of the split luciferase. We tested the ability of the minimal Avian Precursor or WT cattle/Texas polymerases to interact with bovine or human ANP32A or B. We found that the polymerase mutations in cattle/Texas significantly enhanced the ability of the polymerase to interact with both bovine and human ANP32A/B. A stronger interaction was seen with bovine ANP32A compared to ANP32B suggesting that during adaptation to cattle, PB2 M631L and PA K497R have specifically arisen to promote effective binding and use of bovine ANP32A protein.

## Discussion

Our data using reconstructed viral polymerases in host-specific minireplicon and infection assays in cells and explants from different relevant influenza host species show that PB2 M631L is the key adaptive mutation that allowed B3.13 H5N1 high pathogenicity avian influenza virus to efficiently replicate in cattle. PB2 M631L enables the polymerase complex to co-opt bovine ANP32 proteins to support its activity, with a strong preference for bovine ANP32A over ANP32B.This is likely due to bovine ANP32B containing an arginine at position 153, which we have previously shown is responsible for the reduced ability of canine ANP32B to support influenza polymerase^26^. Structurally, in the complex between the influenza polymerase and host ANP32, PB2 residue 631 is located near amino acids 627 and 591 which are well characterised sites that mutate to allow adaptation of AIV polymerase to mammalian ANP32 proteins^26,43^. However, the exact mechanism of how PB2 M631L allows the use of mammalian ANP32s remains unresolved, although our data suggests directly enhanced interaction. Structural studies show that ANP32 bridges the asymmetric dimer of influenza polymerase comprising a replicating subunit and a recruited subunit that will be used to encapsidate the nascent RNA strand. PB2 E627K results in a switch from a negative to a positive charge enhancing the interaction between the encapsidating subunit of the polymerase replication dimer and ANP32 residue E151, while also avoiding the clash between the negatively charged LCAR region of mammalian ANP32s and the replicating polymerase subunit^43,44^. This mechanism also likely applies to adaptive charge switch mutations at residue 591 from Q to R/K^43^. PB2 M631L does not result in an alteration of charge but presumably changes the polymerase structure in this region to better accommodate the interaction between either or both polymerase subunits and ANP32. Recent work from Gu et al. using the cattle virus polymerase confirmed that PB2 M631L increases activity in human cells at both 33°C and 37°C^32^. Previous work has shown that PB2 M631L adapts avian H10N7 viruses to mice^34^ and enables an avian H9N2 virus to use otherwise suboptimal ANP32 proteins^31,45^. These reports, combined with our results in the context of a second 2.3.4.4b genotype and an older non-goose/Guangdong H5N1 suggest that M631L is a general mammalian adaptive mutation for influenza polymerase that is not specific to a particular virus genetic background. However, why this mutation emerged in cattle rather than the more potent PB2 E627K remains unclear. We consistently found when using the minimal Avian Precursor (cattle WT with PB2 M631L + PA K497R removed), the addition of E627K showed lower replication in bovine, but not human systems, than the combined addition of M631L and PA K497R. There was also similar or lower polymerase activity observed in the corresponding minigenome assays. Our previous work has shown that E627K is biased towards emerging in hosts or systems where the virus preferentially uses ANP32B^26^, however in cattle we found that ANP32A better supported the polymerase. E627K has arisen experimentally in cattle infected with a contemporary European H5N1 2.3.4.4b virus^17^.

Following fixation of PB2 M631L in cattle, additional mutations in the polymerase arose in three separate clades. Most cattle sequences contain PA K497R, which increased polymerase activity in the absence of M631L and virus replication in bovine mammary cells. Structurally, PA K497R sits in a loop close to the PB2 M631L and ANP32 interface, though may not be directly interacting with them (Figure 5A). Additionally, PA K497R sits at the FluPol symmetric dimer interface and therefore may modulate the dynamics of dimer formation in the same manner as the mammalian adaptation PA Q556R^46^. Of the mutations defining the other two early cattle virus clades, the combination of PA I13V + E613K or PB2 E677G also showed modest increases in activity, although neither group spread particularly widely. Neither PA I13V or E613K have previously been associated with mammalian adaptation but E613K does fall on the polymerase asymmetric dimer interface (Supplementary Figure S5B) and could be altering polymerase regulation^43^.

Cattle/Texas virus has extremely robust replication in the bovine mammary gland, with very high titres of infectious virus being reported from milk^1,14,17^. However, further mutations that increase polymerase activity in bovine cells are still being reported as the outbreak continues. There have been three separate emergences of PB2 D740N which we show here boosted polymerase activity above the wild-type virus. PB2 E627K has also arisen once in conjunction with M631L in a cluster of cattle and appears to be continuing to expand as of July 2025, consistant with this combination giving a robust enhancement in polymerase activity in combination with M631L in our minireplicon assay, and a replicative advantage in cattle mammary explants. It will be vital to continue to monitor evolution of the polymerase (and other viral genes), and how mutations may impact infectivity and transmissibility of these viruses in cattle and other mammals.

In 2025, USDA reported two further spillovers of H5N1 into cattle in Nevada and Arizona from the D1.1 genotype^20,28^, which was the predominant genotype in wild birds at the time of spillover. Interestingly, neither cluster has PB2 E627K nor PB2 M631L, despite D1.1 PB2 having >98.5% amino acid identity with the B3.13 PB2 (and similarly arising from low pathogenicity avian influenza virus of North American waterfowl). The Nevada cluster has the known mammalian ANP32 adaptive mutation, PB2 D701N^26^, whereas the Arizona cluster has several mutations in the polymerase relative to the closest avian sequences, which have yet to be tested for mammalian activity, such as PB2 R379K and D678N. These independent emergences provide further evidence that E627K does not have a strong selective advantage to the influenza polymerase in cattle, and that there are many different viable pathways for H5N1 polymerase adaption in mammals.

What is the current risk to humans posed by the cattle virus? Our results show that the cattle virus containing mammalian adaptations can replicate better in mammalian cells than the minimal avian-like precursor virus, and is therefore a clear risk to mammals which have consumed infected milk such as cats and raccoons^1^. Thus far, these cattle viruses show inefficient airborne transmissibility in ferret experiments^32,47^ and have not adapted to transmit via respiratory route or to efficiently use human receptors (α-2,6-linked sialic acids) which would be essential for human-to-human transmission^48–51^. In addition, there are also other potential blocks to emergence posed by antiviral host factors such as MxA to which the cattle virus remains susceptible^52^. Nonetheless, frequent exposure of an antigenically novel, high pathogenicity influenza virus that can replicate well in human cells is concerning. Our data suggest that the emerged cattle influenza virus is also capable of efficiently infecting avian and swine cells, further increasing the risk of spillover from cattle to other species. In fact, the mammalian-adapted virus appeared to have no fitness cost in avian cells, explaining the high propensity of this virus to spillover into poultry causing multiple outbreaks, and suggesting such a virus could maintain its mammalian adaptations if it spilt back into wild waterfowl. With each human infection and increased polymerase activity leading to higher levels of replication, there is a danger of further evolution changing viral receptor properties. Additionally, a reassortment event with human seasonal influenza viruses could lead to a novel virus, particularly during the Northern hemisphere winter influenza season.

Recent experimental infections in cows suggest that the ability of the North American B3.13 cattle influenza virus to spread via milk is not unique and that other mammalian viruses are capable of transmitting via this route^17^. Phylogenetic data suggests that this originated from a single spillover event from wild birds to cattle and, despite recent spillovers from D1.1, such spillovers are likely unusual as there is scant evidence of IAV in cattle prior to 2024. Nonetheless, in the absence of an effective control strategy, the high pathogenicity H5N1 virus may now become endemic in US dairy cattle, requiring continuous monitoring even in the absence of overt disease. Moreover, many other clades and strains of H5N1 virus continue to emerge through reassortment, causing zoonotic infections. Urgent development and testing of broadly reactive H5 influenza vaccines for both animals and humans is a priority.

## Methods

### Ethics and biosafety

Virus work was undertaken at either containment level 3 (CL3), SAPO4 (for whole H5N1 reverse genetics-derived viruses) or CL2 (for reverse genetics-derived viruses with HA and NA from the attenuated vaccine strain A/Puerto Rico/8/1934 (PR8) and the remaining internal genes from H5N1 viruses). Viruses carrying H5 HA with a multibasic cleavage site are categorised as specified animal pathogens order (SAPO) 4 and Advisory Committee on Dangerous Pathogens (ACDP) hazard group 3 by United Kingdom regulations. Work with these viruses was undertaken in a licensed CL3/SAPO4 facility of The Pirbright Institute under GMRA (BAG-RA-226). CL2 work with ACDP Hazard Group 2 recombinant influenza viruses was performed at the Roslin Institute under biological risk assessments BARA 1011 and GMRA 1811. All virus and GM risk assessments were approved by the appropriate internal committees, as well as the UK Health and Safety Executive (HSE) and, where necessary, the UK scientific advisory committee for genetic modification (SACGM).

Ethics for the use of the primary human airway epithelial cultures (hNECs) were as described previously^53^. Briefly, donors provided written consent and Ethics approval was given through the Living Airway Biobank, administered through the UCL Great Ormond Street Institute of Child Health (REC reference: 19/NW/0171, IRAS project ID: 261511, Northwest Liverpool East Research Ethics Committee). Nasal brushings were obtained by trained clinicians from adult (30–50 years) donors who reported no respiratory symptoms in the preceding 7 weeks. Brushings were taken from the inferior nasal concha zone using cytological brushes (Scientific Laboratory Supplies, CYT1050). All methods were performed following the relevant guidelines and regulations.

Generation of bovine explants came under approval from the University of Glasgow School of Biodiversity, One Health and Veterinary Medicine (EA26/25).

### Cells

Human Embryonic Kidney 293T (293T), Madin-Darby Canine Kidney (MDCK), MDCK overexpressing chicken ANP32A (MDCK-ggANP32A), chicken immortalised fibroblast cells (DF-1), human adenocarcinoma cells (A549), duck immortalised fibroblasts (CCL-141), human immortalised lung fibroblasts (MRC5), and bovine mammary epithelial cells (MAC-T) were maintained in Dulbecco’s modified Eagle medium (DMEM) supplemented with 10% fetal bovine serum (FBS), and 1% Pen-Strep. MAC-T at Roslin additionally had 2 mM L-glutamine, 5 µg/ml insulin and 1 µg/ml of hydrocortisone. Human lung adenocarcinoma (Calu-3) cells were maintained in DMEM with 10% FCS, 1% Pen-Strep, 2 mM L-glutamine. Swine foetal testes (ST) cells were maintained in Advanced DMEM, 5% FCS, 1% Pen-Strep, 2 mM L-glutamine. Chicken lung epithelial cell (CLEC213^54^, kindly gifted by Dr Sascha Trapp) were cultured in DMEM F12 supplemented with 8% FBS, 2 mM glutamine, 1% Pen-Strep. Triple knockout cells (tKO-Engineered human haploid (eHAP) cells with ANP32A, ANP32B and ANP32E knocked out) were generated as previously described^46^ and maintained in Iscove’s modified Dulbecco’s medium (IMDM), 10% FBS, 1% NEAA, 1% Pen-Strep. All cells were maintained at 37 °C and 5% CO_2_.

Primary human nasal epithelial cultures (hNECs) were generated as described previously^53^. Briefly, basal epithelial cells from nasal brushings were expanded on mitotically inactivated 3T3-J2 fibroblasts to passage 1 and cryopreserved. Cells were thawed as required, expanded with 3T3-J2 fibroblasts to passage 2, and seeded onto collagen I–coated, semi-permeable membrane supports (3 × 10^5 cells/6.5 mm, 0.4 µm pore size, Transwell; Corning). Differentiation was performed under air–liquid interface conditions in PneumaCult™-ALI medium for 4 weeks.

Bovine mammary tissue was collected at Sandyford Abattoir from health lactating cattle. Excess tissue was removed, teat and gland cisterns were dissected and transferred to transport media comprising of chilled DMEM supplemented with 100 μg/ml Penicillin/ 100 μg/ml streptomycin and 5 μg/ml fungizone. Every 45 minutes, tissues were transferred to 500ml of fresh transport media. Inside class II biological safety cabinet, explants were made using a 5mm biopsy punch. Biopsies were added to 24 well plates with 500ul of transport media supplemented with 10% FBS and incubated for 24h at 37°C, 5% CO*_2_*before infection.

### Plasmids

Reverse genetics plasmids of the viruses used in this study were produced as previously described (Table 1)^51,55,56^. Mutants were generated by site directed mutagenesis. pCAGGS expression plasmids of the polymerase subunits were subcloned from reverse genetics plasmids.

**Table 1.** Virus strains, aliases, accessions and uses in this study.

| Virus strain name | Alias | GISAID ID | Whole virus, 2:6 or minireplicon |
| --- | --- | --- | --- |
| A/Puerto Rico/8/1934 | PR8 |  | 2:6 HA and NA only |
| A/dairy cattle/Texas/24-008749-001-original/2024 (H5N1) | cattle/Texas | EPI_ISL_19094762 | All |
| A/chicken/England/053052/2021(H5N1) | AIV07 | EPI_ISL_9012457 | minireplicon only |
| A/turkey/England/50-92/1991(H5N1) | 50-92 | EPI_ISL_5888 | minireplicon only |
| A/England/195/2009 | Eng195 | EPI_ISL_29994 | Whole virus and minireplicon only |

Construction of the minireplicon luciferase reporter plasmid for use in bovine cells was performed using a previous strategy used for an equivalent swine reporter, pSPOM2-Firefly^39^ with swine polI promoter replaced with 428bp bovine polI promoter from the bovine genome, bosTau8 chrUn_GJ059828v1 (1165-1592)^57^. Plasmid map available at github.com/Flu1/bovinePolI. *Bos taurus Anp32A* (NCBI reference sequence NM_001195019.1) and *Anp32B (NM_001035074.1)* were ordered as synthetic GeneArt constructs from ThermoFisher Scientific and subcloned into pCAGGS as described previously^58^.

### Viruses

Viruses were rescued using 8 plasmid pHW2000 (or derivates) sets containing each genome segment from the named virus. HEK 293T cells were transfected with a mix containing 250 ng of each plasmid using lipofectamine 2000 according to the manufacturer’s instructions. Media was replaced 24 h post-transfection with serum-free DMEM with 0.14% bovine serum albumin and 1 μg/ml of L-(tosylamido-2-phenyl) ethyl chloromethyl ketone (TPCK)-treated trypsin. Virus-containing supernatants were collected after 48 h, clarified by centrifugation and used to infect MDCK-ggANP32A cells or inoculate 10-day-old embryonated hen’s eggs. Infected MDCK-*Gg*ANP32A were monitored twice daily and supernatant was harvested, clarified, aliquoted and stored when clear cytopathic effect was observed (generally between 48 and 72 hpi).Eggs were incubated for 48 h before humane euthanization at 4°C, after which virus-containing allantoic fluid was collected, clarified and stored at −80°C before titration by plaque assay.

### Phylogenetics

Dairy cattle sequences submitted prior to 1^st^ December 2024, avian H5N1 sequences from the USA submitted between 1^st^ September 2023-31^st^ March 2024 and A/Texas/37/2024 were downloaded from GISAID. Sequences from isolates which contained PB2, PB1, PA, NP, HA, NA, NS1, NS2/NEP, M1 and M2 segments were concatenated. Sequences were aligned using MAFFT v7.490 trimmed using pytrimal with a gap threshold of 0.85 to ensure correct residue numbering (total length = 4466 amino acids). Duplicate sequences and sequences with > 30 ambiguous residues or gaps were removed. A phylogenetic tree from the cleaned alignment (n=902) was made in IQTree2 running automatic ModelFinder and the FLAVI+F+I+R2 model^59,60^. Data manipulation and visualisation was carried out using Python 3.12.2 and R 4.4.1. Code and list of GISAID accessions can be found at https://github.com/fernandocapelastegui/Cattle-H5N1-polymerase-adaptation.

### Minireplicon assay

293T, MAC-T, or eHAP tKO cells were transfected in 24-well plates with Lipofectamine 3000 transfection reagent (Thermo Fisher) with the following amounts of pCAGGS expression plasmids: 40 ng PB2, 40 ng PB1, 20 ng PA, 80 ng NP, 40 ng Renilla luciferase, 80 ng species-specific (human, bovine, swine or chicken) polI vRNA Firefly luciferase and 40 ng ANP32 or empty pCAGGS (where appropriate). DF-1 and ST cells were transfected in 12 well plates using quadruple the plasmid/transfection reagent mixes to those used for the other cells in 24-well plates. Cells were lysed 24h post-transfection using passive lysis buffer (Promega) and polymerase activity measured using the Dual-luciferase Reporter assay system (Promega) and a FLUOstar Omega plate reader (BMG Labtech). Assays in MAC-T cells were read using Dual-Glo® Luciferase Assay System (Promega) on an Infinite MPlex plate reader (Tecan). Firefly luciferase signal has been normalised to Renilla luciferase signal to give relative luminescence units (RLU).

### Virus replication kinetics

Confluent monolayers of CLEC213, CCL-141, A549, MRC5, MAC-T, BAT II, and Calu-3 cells were washed once with PBS and infected at multiplicity of infection (MOI) 0.01 with virus diluted in serum-free medium for 1 h at 37 °C. Inoculum was replaced with serum-free medium supplemented with 0.14% BSA and 1 µg/ml TPCK-treated bovine pancreas trypsin. Calu-3 and MAC-T experiments performed with whole H5N1 virus excluded TPCK trypsin. Supernatants were collected at various times post infection and stored at −80°C until infectious titres were determined by plaque assay.

Infection of primary human airway cltures was performed as follows. Before infection cells were washed with Dulbecco’s phosphate-buffered saline supplemented with calcium and magnesium (DPBS++) to remove mucus and debris. Cells were infected with 200 µl of virus in DPBS++ at an MOI of 0.01 and incubated at 37°C for 1 h. Inoculum was removed, and cells were washed twice with DPBS++. Time points were taken by adding 200 µl of DPBS++ and incubating for 10 mins and 37°C before removal and titration. The time course was performed at 37 °C, 5% CO_2_. Time points were taken at 24-, 48-, 72-and 96-hours post-infection. Infectious titres were determined by plaque assay on MDCKs.

Infection of explants was performed as follows: individual explants were transferred to 96-well plates and infected with 5000 PFU of virus in 100 µl of serum free DMEM for 2h at 37 °C. Explants were transferred to clean 24-well plates, washed 3x with warm PBS and overlayed with 1 ml of DMEM supplemented with 0.14% fraction V BSA and 1 µg/ml TPCK. Time points were collected at the indicated times post infection.

### Split Luciferase assays

Split luciferase assays were performed in 293T cells seeded into 48-well plates. 40 ng of PB2, PA, and PB1 tagged with the N-terminus of *Gaussia* luciferase (Gluc1) on its C-terminus (after a GGSGG linker), were co-transfected with ANP32 tagged with the C-terminus of *Gaussia* luciferase (Gluc2) on its C-terminus (after a GGSGG linker), using lipofectamine 3000 as per the manufacturer’s instructions. 24 hours post-transfection, cells were lysed in 60 µl of *Renilla* lysis buffer (Promega), and luciferase activity was measured using a *Renilla* luciferase kit (Promega) on a VANTAstar plate reader (BMG Labtech). Normalised luminescence ratios (NLRs) were calculated by diving the values of the tagged PB1 and ANP32 wells by the sum of the mean of the control wells, which contained i) untagged PB1 and free Gluc1, and ii) untagged ANP32 and free Gluc2, as described elsewhere^24^.

### Immunoblotting

Transfected cells were lysed in radioimmunoprecipitation (RIPA) buffer (150mM NaCl, 50mM TRIS, 0.1% SDS, 1% NP-40, 0.5% sodium deoxycholate, pH 7.4) supplemented with an EDTA-free protease inhibitor mini tablet (Thermo Fisher), incubated on ice for 10 minutes, then centrifuged for 10 minutes to remove precipitate. Lysates were then diluted in 4 x Laemmli buffer (Bio-rad) and loaded onto 4-20% gradient gels to perform SDS-PAGE. Proteins were transferred onto a PVDF membrane by semi-dry transfer. Membranes were blocked in TBS/5% milk for 1 hour at room temperature and incubated overnight at 4°C with the following primary antibodies: mouse anti-FLAG (Sigma, F1804; 1:250 dilution), rabbit anti-PB2 (GeneTex, GTX125926; 1:500 dilution), rabbit anti-PB1 (Genetex, GTX125923, 1:250 dilution), rabbit anti-PA (Genetex, GTX118991; 1:500 dilution), mouse anti-α-Tubulin (Abcam, ab7291; 1:1250 dilution) rabbit anti-*Gaussia* luciferase (Invitrogen, PIPA1181; 1:1000). Following four 10-minute TBS/1% Tween20 (TBS-T) washes, membranes were incubated with the following near-infrared fluorescent secondary antibodies: goat anti-mouse IgG Alexa FluorRTM 680 (abcam; ab175775; 1:10,000 dilution), goat anti-rabbit IRDye 800CW (LI-COR, 926-32211; 1:10,000 dilution) for 45 minutes at room temperature. After a further four TBST washes, and one TBS wash, membranes were imaged using an Odyssey DLx (Li-Cor Biosciences) using the Image Studio Lite software.

### Virus whole genome sequencing

Viral RNA was extracted from the virus inoculums used in hNECs, and the 72-hour time point using the QIAamp viral RNA extraction kit (Qiagen) according to the manufacturer’s instructions. Viral cDNA was synthesised using the Veso cDNA synthesis kit (Thermo) with a mix of UTR-specific primers (TATTCGTCTCACCCAGCAAAAGCAGG and TATTCGTCTCACCCAGCGAAAGCAGG) with cycling conditions of 42°C for 75 min, 95°C for 2 min. The Optil-F1 (GTTACGCGCCAGCAAAAGCAGG), Optil-F2 (GTTACGCGCCAGCGAAAGCAGG) and Optil-R1 (GTTACGCGCCAGTAGAAACAAGG) primers were used in a PCR from cDNA with PFU Ultra II DNA polymerase (Agilent). Cycling conditions were 55°C for 2 minutes, 98°C for 2 minutes. 5 x cycles of 94°C for 30s, 44°C 30s, 68°C 3m30s, 35x cycles of 94°C for 30s, 57°C for 30s, 68°C 3m30s, a final extension step of 68°C for 10m followed by a holding step of 4°C. The PCR products were then purified using the Agencourt AMPure XP kit per manufacturer’s instructions. Purified amplicons were quantified by Qubit (Thermo) and diluted in nuclease-free water to a final concentration of 0.2ng/µl. 1 ng total of input nucleic acid was processed using the Illumina Nextera XT kit on a Hamilton NGStar. The resulting libraries were confirmed using the Tapestation 4200 *Agilent) and bead-normalised according to the manufacturer’s specifications before being pooled and diluted to 12.5 pM final concentration with 1% 12.5 pM PhiX (Illumina). The resulting pool was then run on an Illumina MiSeq using a 2 x 250 v2 MiSeq reagent cartridge and flow cell.

Raw fastq sequences were then imported into Geneious Prime software and trimmed to remove primers and aligned to the whole genome reference sequence (A/dairy cattle/Texas/24-008749-001-original/2024 (H5N1)).

## Acknowledgements

MRC-5 cells were a kind gift from Dr Finn Grey of the Roslin Institute. Chicken lung cells (CLEC213^54^) were kindly gifted by Dr Sascha Trapp of INRA Centre Val de Loire. CCL-141 cells were a kind gift from Dr Leah Golding of University of Nottingham. MDCK-ggANP32 cells were a kind gift from Professor Massimo Palmarini of the Centre for Virological Research, Glasgow. The authors would like to thank Dr Martha Nelson for feedback on the manuscript and Dr Giulia Gallo for useful suggestions on data visualisation. The authors thank all researchers who have shared genome data openly via the Global Initiative on Sharing All Influenza Data (GISAID).

V.D. and F.C. were supported by Academy of Medical Sciences Springboard Grant 1049 and Royal Society Grant 231225 to D.H.G., J.L.Q., T.M, M.N.J.W., C.M.S., N.P., R.M.P., M.D.B., J.Y., S.R., P.M., I.H.B, M.I., P.D., W.B. T.P.P. and D.H.G are supported by the UK Medical Research Council/Department for Environment, Food and Rural Affairs (Defra, UK) FluTrailMap-One Health consortium [MR/Y03368X/1]. J.L.Q., R.M.P., M.D.B., I.H.B., M.I., P.D., W.S.B., and T.P.P. are also supported by the Biotechnology and Biological Sciences Research Council (BBSRC)/DEFRA ‘FluTrailMap’ consortium [BB/Y007298/1]. J.L.Q., J.Y., I.H.B., M. I., P.D., W.S.B., R.M.P, and T.P P. were supported by the BBSRC/Defra “FluMAP” consortium (BB/X006123/1). S.R., G.F., A.M., I.H.B., M.I. and T.P.P. are funded by the BBSRC via the Pirbright Institute’s Strategic Programme Grants (ISPGs) [BBS/E/PI/230002A; BBS/E/PI/230002B], BBSRC National Bioscience Research Infrastructure: High Containment and Low Containment Services and Science Platforms grants [BBS/E/PI/23NB0004, BBS/E/PI/23NB0003]. ES was supported by Medical Research Council (MRC) programme grant MR/X008312/1. P.D. and R.M.P. acknowledge BBSRC Strategic Programme Grant funding to the Roslin Institute (BBS/E/RL/230002C). P.R.M. is supported by BBSRC [BB/V004697/1]. R.M.P is supported by a Chancellor’s Fellowship from the University of Edinburgh. P.D. also acknowledges BBSRC support from the Evolution & Ecology of Infectious Diseases programme (BB/V011286/1), MRC support from a ONE Health epidemic preparedness award (MR/Y015045/1) and Defra award FluSwitch (SE2223).

## Credit Contributions

VD – investigation, resources, visualisation

JLQ – investigation, resources

SR – investigation, Writing – review & editing

NP – investigation, Writing – review & editing

MDB – investigation, resources, Writing – review & editing

JY – investigation, resources, Writing – review & editing

FC – data curation, investigation, software, Writing – review & editing

TM - resources

K-MC – resources

JA – resources

MNJW – resources

CM - resources

GF – methodology, investigation, supervision

AM – methodology, investigation

ES – resources, Writing – review & editing

CMSheppard – investigation, supervision, Writing – review & editing

IHB – funding acquisition, supervision, Writing – review & editing

PRM - funding acquisition, resources, supervision

CMSmith - funding acquisition, resources, supervision

MI - funding acquisition, resources, supervision

PD - funding acquisition, methodology, supervision, Writing – review & editing

WSB – conceptualisation, funding acquisition, methodology, supervision, Writing – review & editing

RMP – investigation, methodology, project administration, supervision, validation

TPP – conceptualisation, formal analysis, investigation, methodology, project administration, supervision, validation, visualisation, Writing – original draft

DHG – conceptualisation, data curation, formal analysis, funding acquisition, investigation, methodology, project administration, software, supervision, visualisation, Writing – original draft

