## Supplementary Figures for "Polymerase mutations underlie early adaptation of H5N1 influenza virus to dairy cattle and other mammals"

Supplmentary Figures

Supplementary Figure S1

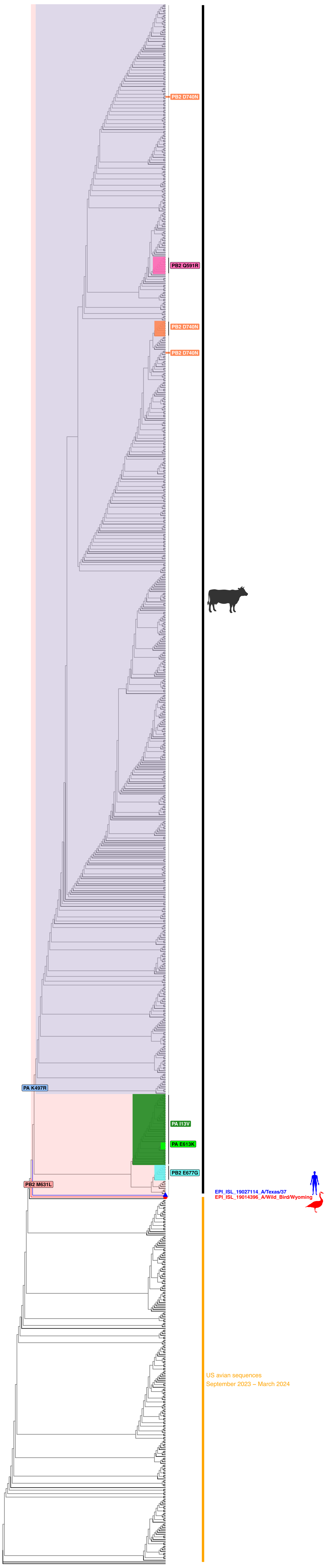

**Supplementary Figure S1. Phylogenetic analysis of bovine H5N1 viruses with key polymerase amino acid changes annotated.** Influenza sequences in the cattle outbreak, USA  avian H5N1 sequences from 1st September 2023 – 31st March 2024 and A/Texas/37/2024 were concatenated and duplicates removed. The phylogenetic tree was created in IQTree2. Key polymerase mutations defining clades are highlighted.

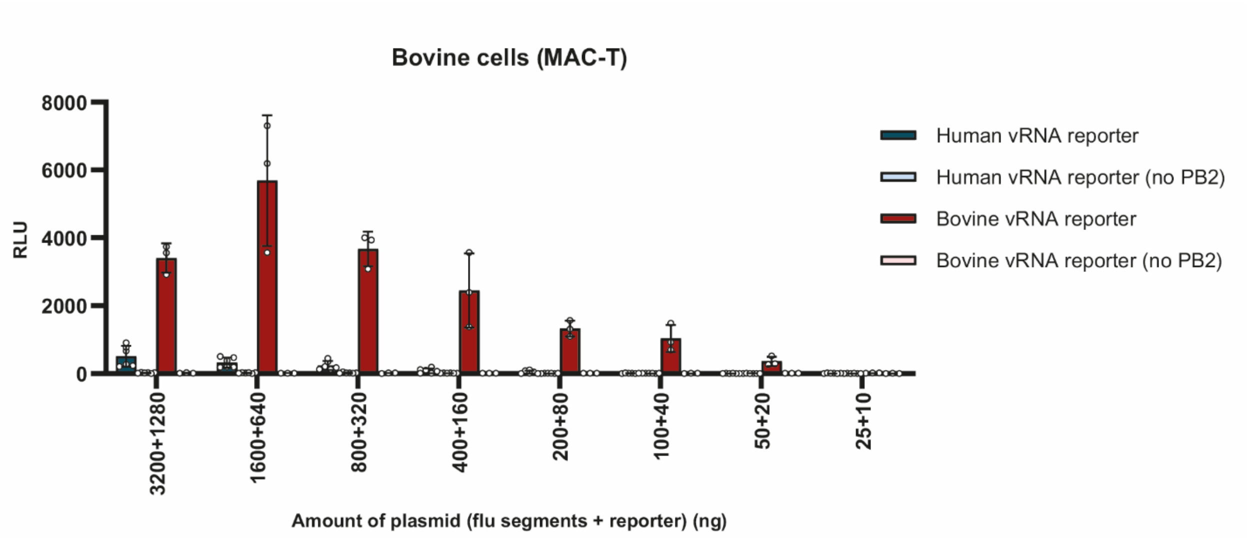


**Supplementary Figure S2. Generation and validation of a bovine cell-specific minireplicon reporter.** Validation of bovine-cell specific reporter vs human-cell specific reporter in bovine MAC-T cells. Data plotted as mean ± SD. Minireplicon assay performed with either whole PR8 polymerase + NP, or PR8 polymerase + NP missing PB2.

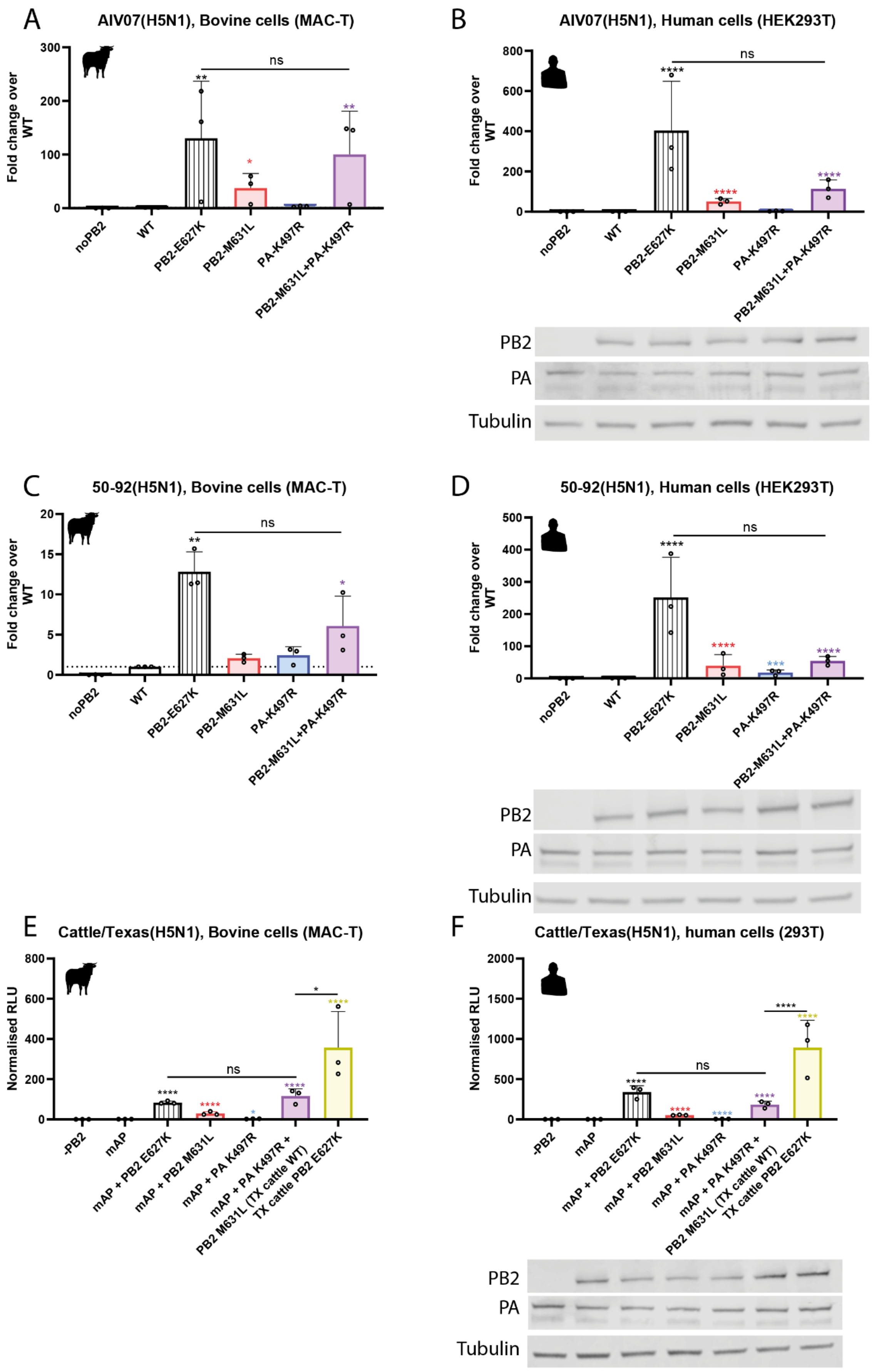
A
AIV07(H5N1), Bovine cells (MAC-T)
Fold change over WT
300
200
100
0
ns
**
*
**
noPB2
WT
PB2-E627K
PB2-M631L
PA-K497R
PB2-M631L+PA-K497R
B
AIV07(H5N1), Human cells (HEK293T)
Fold change over WT
800
600
400
200
0
ns
****
****
****
noPB2
WT
PB2-E627K
PB2-M631L
PA-K497R
PB2-M631L+PA-K497R
PB2
PA
Tubulin
C
50-92(H5N1), Bovine cells (MAC-T)
Fold change over WT
20
15
10
5
0
ns
**
*
noPB2
WT
PB2-E627K
PB2-M631L
PA-K497R
PB2-M631L+PA-K497R
D
50-92(H5N1), Human cells (HEK293T)
Fold change over WT
500
400
300
200
100
0
ns
****
****
***
****
noPB2
WT
PB2-E627K
PB2-M631L
PA-K497R
PB2-M631L+PA-K497R
PB2
PA
Tubulin
E
Cattle/Texas(H5N1), Bovine cells (MAC-T)
Normalised RLU
800
600
400
200
0
ns
*
****
****
****
*
****
-PB2
mAP
mAP + PB2 E627K
mAP + PB2 M631L
mAP + PA K497R
mAP + PA K497R + PB2 M631L (TX cattle WT)
TX cattle PB2 E627K
F
Cattle/Texas(H5N1), human cells (293T)
Normalised RLU
2000
1500
1000
500
0
ns
****
****
****
****
****
****
-PB2
mAP
mAP + PB2 E627K
mAP + PB2 M631L
mAP + PA K497R
mAP + PA K497R + PB2 M631L (TX cattle WT)
TX cattle PB2 E627K
PB2
PA
Tubulin

**Supplementary Figure S3. Polymerase mutations found in the major cattle clade also enhance mammalian cell polymerase activity in an unrelated genetic background.** Minigenome in (A,C,E) bovine MAC-T cells (B,D,F) human HEK293T, with mutant versions of a contemporary 2.3.4.4b UK H5N1 strain, A/chicken/England/053052/2021(H5N1) a.k.a. AIV07 (A,B) and the historic, non-goose/Guangdong H5N1 strain, A/turkey/England/50-92/1991(H5N1)(C,D) or of Cattle/Texas (E,F). Data normalised to WT. (B,D,F) Matched western blots showing expression of PB2 and PA. Data throughout plotted as the mean of N = 3 independent biological repeats. Data throughout plotted as mean + SD. Statistics throughout performed by one-way ANOVA with multiple comparisons comparing all data points, only comparisons between WT and mutants, and between PB2-E627K vs PB2-M631L+PA K497R marked on the figures. Log-normality determined by Shapiro-Wilk test and QQ plot. Significance shown by asterisks indicating: *, 0.05 ≥ P > 0.01; **, 0.01 ≥ P > 0.001; ***, 0.001 ≥ P > 0.0001; ****, P ≤ 0.0001.

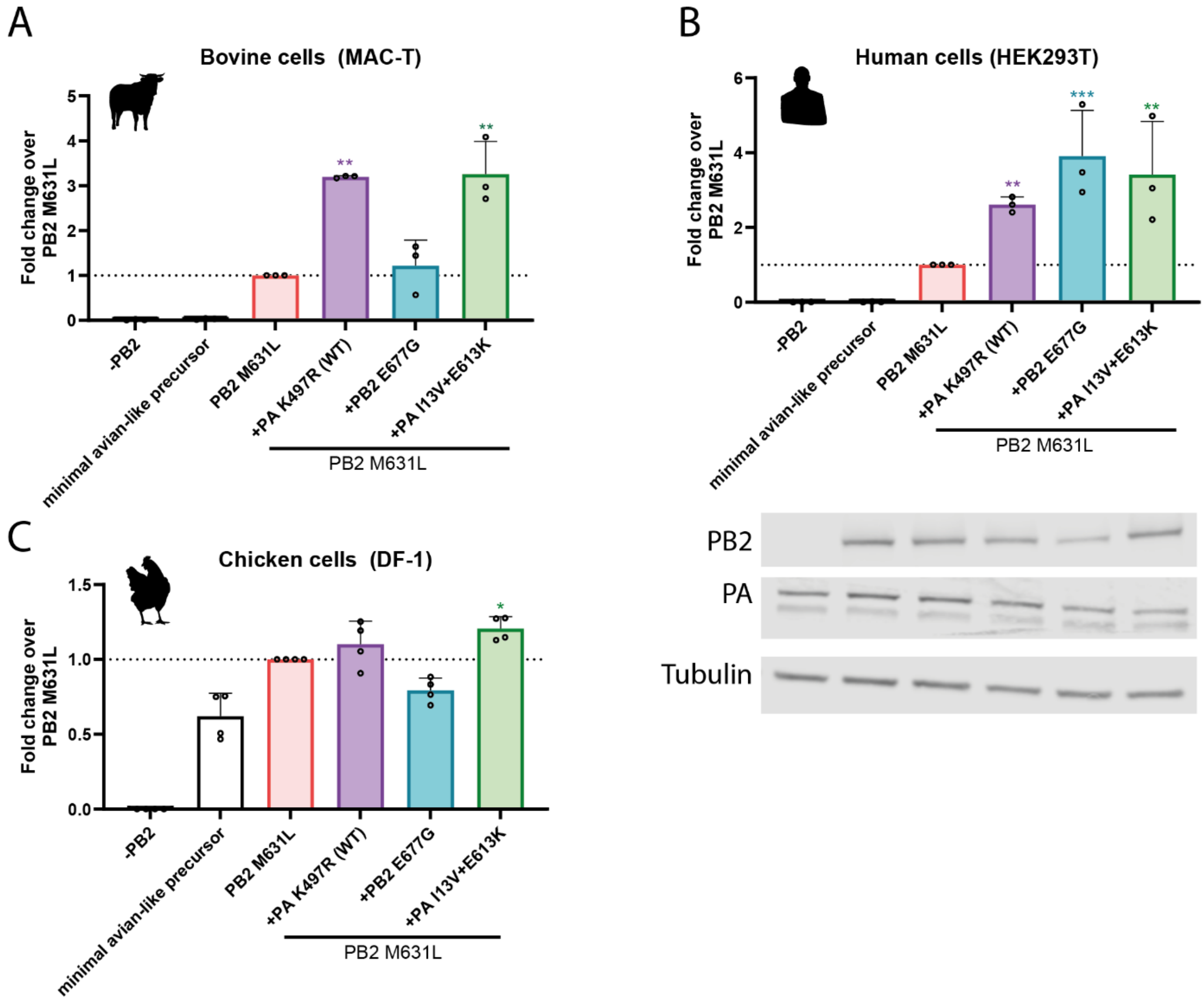

**Supplementary Figure S4. Polymerase mutations on minor cattle branches also enhance polymerase activity in mammalian cells.** Minigenome in (A) bovine MAC-T cells (B) human HEK293T, (C) chicken DF-1 with mutant versions of cattle Texas. Data normalised to PB2 M631L. B) Expression of PB2 and PA from a matched western blot. Data throughout plotted as the mean of three independent biological repeats. Data throughout plotted as mean + SD. Statistics throughout performed by one-way ANOVA with multiple comparisons against PB2 M631L using log-transformed data. Log-normality determined by Shapiro-Wilk test and QQ plot. Significance shown by asterisks indicating: *, 0.05 ≥ P > 0.01; **, 0.01 ≥ P > 0.001; ***, 0.001 ≥ P > 0.0001; ****, P ≤ 0.0001.

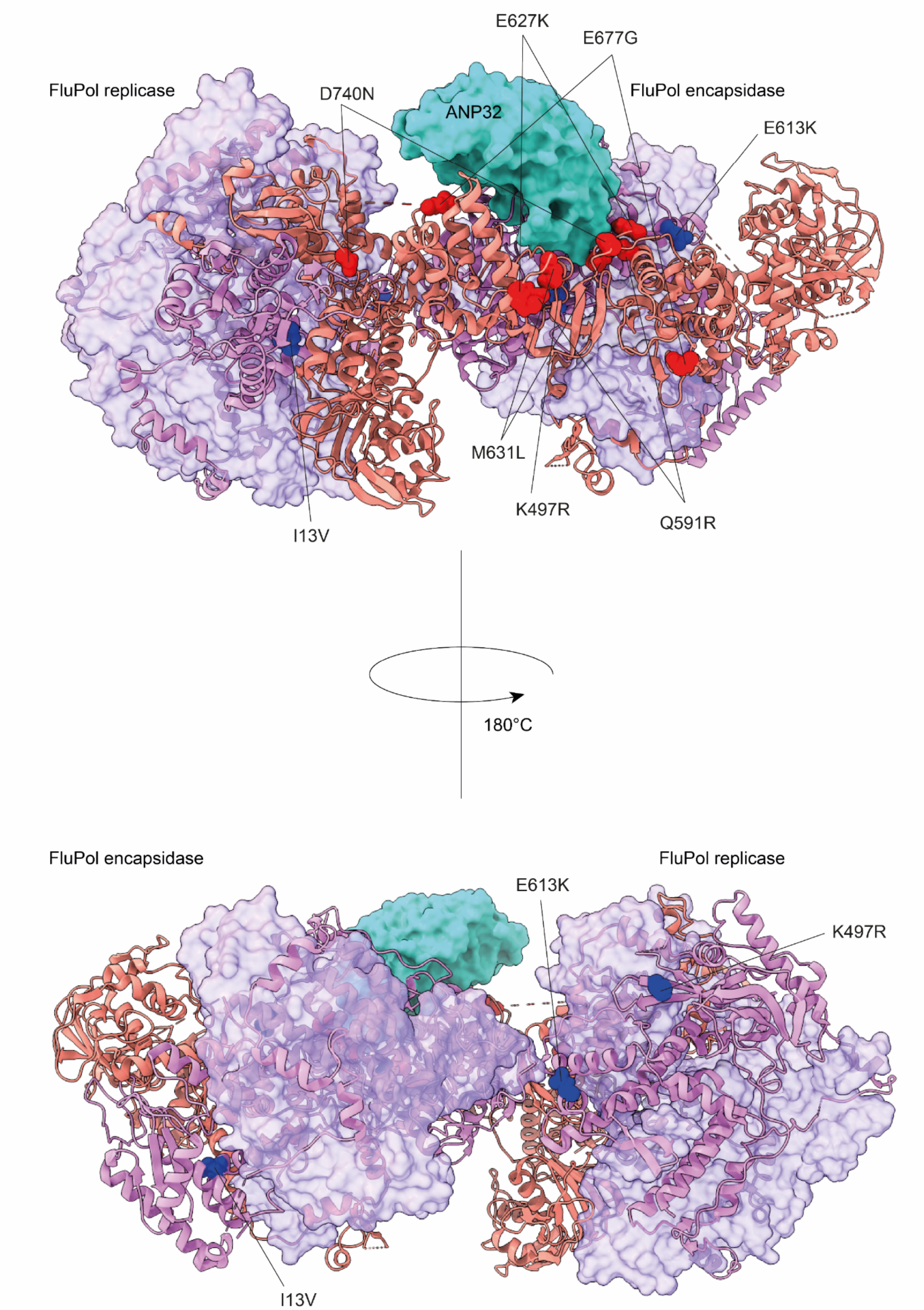


**Supplementary Figure S5: Extended structural mapping of cattle mutations onto the influenza virus polymerase.** H5N1 influenza A virus asymmetric polymerase dimer (PDB: 8R1J) showing human ANP32B in cyan, PB1 in purple, PB2 in salmon and PA as ribbon representation in violet. The H5N1 cattle PA substitutions sites (13, 497, 613) are highlighted in blue and PB2 substitutions (591, 627, 631, 677, 740) are highlighted in red.
